# Metabolic shifts in the heart of hypothyroid mice — insights from untargeted metabolomics

**DOI:** 10.1101/2025.01.16.633354

**Authors:** Heng Guan, Yujie Yang, Wenping Xiao, Sha Wan, Fang Liu

**Affiliations:** Department of Anatomy, College of Basic Medicine, Guilin Medical University, Guilin, China; Department of Surgical Comprehensive Ward, Longgang District People’s Hospital of Shenzhen, Shenzhen, China; Taikang Center for Life and Medical Sciences, Wuhan University, Wuhan, China; Center of Diabetic Systems Medicine, Guangxi Key Laboratory of Excellence, Guilin Medical University, Guilin, China

**Keywords:** Hypothyroidism, Cardiac atrophy, Untargeted metabolomics, Fatty acid oxidation, Phospholipid metabolism

## Abstract

**Introduction:** Hypothyroidism is a prevalent disorder that affect all systems of body, including the cardiovascular system and control of cardiac function. However, much remains unknown about the metabolic pathways and disorders in the heart induced by hypothyroidism, as well as the metabolic mechanisms involved.

**Objectives:** This work aims to investigate the impacts of hypothyroidism on the heart and explore its underlying mechanisms through untargeted metabolomics analysis.

**Methods:** An experimental model of hypothyroidism was established by giving water containing 0.1% methimazole and 1% potassium perchlorate to 12-week-old C57BL/6J mice for six consecutive weeks.

**Results:** After treatment, hypothyroidism significantly reduced mice’s heart weight and heart rate. Metabolomics analysis revealed that the different metabolites (DEMs) primarily concentrated in Phospholipids, Lipids or Lipids-like, Carbohydrates and Carbohydrate conjugates. These specific metabolites were mainly enriched in pathways related to lipid metabolism. The hearts of hypothyroid mice exhibited significantly lower levels of lipoyl-carnitine compared to controls, along with markedly higher levels of glycolysis and pentose phosphate pathway (PPP) intermediates. Furthermore, carnitine palmitoyltransferase 1B (CPT1B) and carnitine palmitoyltransferase 2 (CPT2) expressions were significantly decreased in the hypothyroid mouse heart, while lactate dehydrogenase A (LDHA) expressions was significantly increased, according to the immunoblotting assay.

**Conclusions:** Research indicates that hypothyroidism triggers disruptions in both cardiac energy metabolism and phospholipid metabolism in adult mice, which in turn results in myocardial atrophy. These findings could pave the way for novel therapeutic strategies aimed at treating hypothyroidism-induced myocardial atrophy.

## 1 Introduction

Hypothyroidism, a disease with multiple etiologies, is characterized by thyroid dysfunction or thyroid hormone (TH) resistance, resulting in a systemic hypometabolic syndrome. Almost all tissues and organs in the body are regulated by TH, alterations in TH levels may impact each physiological system, with the cardiovascular system being mostly affected^[1,2]^. TH has a central regulatory role in the cardiovascular system, especially in the heart, and significant changes in cardiac function are often closely linked to fluctuations in plasma or tissue levels of TH^[3]^. Furthermore, the actions of thyroid-stimulating hormone (TSH) can extend beyond the thyroid gland. Therefore, greater levels of TSH must be considered while investigating the consequences of hypothyroidism on the heart^[4]^.

Previous studies have identified a physiological phenomenon in which the expression of sarcoplasmic/endoplasmic reticulum Ca^2+^ ATPase 2a in the hearts of laboratory rodents with hypothyroidism is considerably reduced by low TH levels or high TSH levels^[5–7]^. This inhibition led to a decrease in the efficiency of Ca^2+^ uptake, which triggered the impairment of myocardial systolic and diastolic function. Moreover, hyperpolarization-activated cyclic nucleotide-gated channel expression was decreased by increased TSH levels in hypothyroid animals, which in turn regulated cardiac function and caused ventricular muscle atrophy and cardiac contractile dysfunction ^[8]^. The earlier experiments demonstrate the complex processes by which hypothyroidism impacts heart health. Similarly, our hypothyroid mice displayed atrophied and damaged myocardium along with diminished cardiac function.

Metabolomics has the unique ability to quantitatively capture the comprehensive and dynamic metabolic response that living systems exhibit in the face of pathological stimuli or disturbances. It focuses on tiny modifications in endogenous molecule composition, which function as a mirror reflecting the dynamic responses of organisms to disease or environmental changes^[9]^. Metabolomics opens a window into the inner workings of organisms and their sensitive reactions to environmental changes by closely examining these fluctuations at the molecular level.

Cardiomyocytes can respond precisely to changes in endogenous metabolites caused by hypothyroidism. Describe these responses in detail by metabolomics, which provides a comprehensive metabolic understanding of the mechanisms behind the onset and development of cardiac disease. Consequently, 14 heart samples were extracted from normal and hypothyroid mice in this study to reveal a list of metabolites associated with hypothyroidism. Next, we performed untargeted metabolomics testing on all samples, screened for DEMs, and performed Kyoto Encyclopedia of Genes and Genomes (KEGG) pathway enrichment analysis. We speculated that this study provides new ideas for exploring the mechanisms by which hypothyroidism leads to the development and progression of myocardial atrophy.

## 2 Materials and Methods

### 2.1 Animal and treatment

All experimental procedures were approved by the Institutional Animal Care and Use Committee of Guilin Medical University. The male C57BL/6J mice used in this study were acquired from Hunan Slake Jinda Laboratory Animal Co. Ltd. They were raised in SPF-class animal houses under controlled conditions of lighting (12-hour light/dark cycle), ambient temperature (20-25°C), and humidity (50%-55%). Food and water were freely accessible to mice. The mice of 12 weeks were randomly divided into control and hypothyroid groups (30 mice in each). Based on previous result^[10]^, antithyroid drugs were consumed to mice as a hypothyroid model. Briefly, mice in the hypothyroid group were continuously given water supplemented with 0.1% (wt/vol) methimazole (Sigma-Aldrich, M8506-25G) and 1% (wt/vol) potassium perchlorate (Aladdin, P103994) for six weeks. Body weights were recorded every week to track their condition of growth and development.

### 2.2 Electrocardiogram (ECG)

The mice were anesthetized with of 2% pentobarbital sodium (0.01ml per 1g body weight). In following the guidelines for electrocardiograph (TECHMAN, BL-420S), four electrodes were then placed subcutaneously on each of the four limbs. This was done with the electrode tips pointing in the direction of the heart to get the most precise ECG readings. The ECG was recorded when the monitor showed a relatively stable heart rate.

### 2.3 Sample collection

Blood samples were collected. Then the heart was quickly removed from the thoracic cavity, and the blood was washed out with cold saline before the water was blotted out on absorbent paper and weighed. After being weighed, the several that were used for the metabolomics of the heart were immediately quenched in liquid nitrogen and kept at -80°C, while the other several were kept at -80°C directly. The thyroid gland was then extracted and preserved in 4% paraformaldehyde. At last, the tibia of the right leg was taken out and its length measured.

### 2.4 Mouse serum ELISA assay

Blood samples were left at room temperature for two hours. The serum was collected after centrifugation at 1200 rpm for 10 min at 4°C, then immediately stored at -80°C. Commercial ELISA kits (CUSABIO, CSB-E05116m, CSB-E05083m, and CSBE05086m) were used to measure TSH, TT4, and TT3 concentrations. The concentration of the sample was calculated by drawing a standard curve based on the concentration and OD value of the standard.

### 2.5 Untargeted metabolomics

#### 2.5.1 Sample preparation

Add 1 milliliter of cool 90% methanol to 80 mg of heart. The MP homogenizer (24×2, 6.0M/S, 60s, twice) was used to homogenize the lysate. The mixture was centrifuged for 20 minutes at 14000g and 4 °C after the homogenate was sonicated at low temperature (30 minutes once or twice). A vacuum centrifuge was used to dry the supernatant. The samples were redissolved in 100 μL of acetonitrile (Merck,1499230-935)/water (1:1, v/v) solvent for LC-MS analysis.

#### 2.5.2 LC-MS/MS Analysis

A quadrupole UHPLC (1290 Infinity LC, Agilent Technologies) coupled to a quadrupole time-of-flight (AB Sciex TripleTOF 6600) was used for the analysis. Samples were analyzed using a 2.1 mm × 100 mm ACQUIY UPLC BEH 1.7 µm column (Waters, Ireland) for HILIC separation. The mobile phase contains A=25 mM ammonium acetate, 25 mM ammonium acetate, and 25 mM ammonium hydroxide in water, and B=acetonitrile in both the ESI positive (POS) and negative (NEG) modes. The gradient initially at 85% B for 1 min was linearly decreased to 65% in 11 min, then quickly dropped to 40% for 4 min, and finally returned to 85% in 0.1 min, followed by a 5-min re-equilibration.

A 2.1 mm × 100 mm ACQUIY UPLC HSS T3 1.8 µm column (Waters, Ireland) was employed for RPLC separation. The mobile phase contained A= water with 0.1% formic acid and B= acetonitrile with 0.1% formic acid in ESI POS mode, and the mobile phase contained A=0.5 mM ammonium fluoride in water and B= acetonitrile in ESI NEG mode. The gradient was 1% B for 1.5 min, increased linearly to 99% in 11.5 min, held for 3.5 min, then reduced to 1% in 0.1 min, followed by a 3.4 min re-equilibration. The gradients were at a flow rate of 0.3mL/min, and the column temperatures were kept constant at 25 ℃. Each sample was injected as a 2 µL aliquot.

The ESI source parameters were configured with Ion Source Gas 1 and 2 at 60, curtain gas at 30, source temperature at 600 ℃, and IonSpray Voltage at ± 5500 V. In MS-only acquisition, the m/z range was set from 60 to 1000 Da, with a TOF MS scan accumulation time of 0.20 seconds per spectrum. In auto MS/MS acquisition, the m/z range was expanded to 25-1000 Da, and the accumulation time for the production scan was set at 0.05 s/spectra. Utilizing information dependent acquisition in high sensitivity mode. The collision energy was fixed at 35 V with ±15eV; while declustering potential was set to 60V for POS ions modes and -60V for NEG ions modes. Isotopes within a 4Da range were excluded, and the system was configured to monitor up to 10 candidate ions per cycle.

#### 2.5.3 Data processing

ProteoWizard MSConvert was used to convert the raw MS data to MzXML files, which were then imported into the open-source XCMS software. Peak picking parameters included centWave m/z at 10 ppm, peakwidth c (10, 60), and prefilter c (10, 100). For peak grouping, the parameters were bw = 5, mzwid = 0.025, and minfrac = 0.5. Collection of Algorithms of MEtabolite pRofile Annotation was used for the annotation of isotopes and adducts. Only variables with more than 50% of the nonzero measurement values in at least one group were retained in the extracted ion features. Metabolite compound identification was performed by comparing accuracy m/z value (<10 ppm), and MS/MS spectra with an in-house database established with available authentic standards. The K-Nearest Neighbor method filled in the missing data and the extreme values were deleted. Lastly, to guarantee the parallelism between samples and metabolites, the total peak area of the data was normalized.

### 2.6 Western blot analysis

The hearts were completely homogenized in RIPA Lysis Buffer (Beyotime, P0013B) and PMSF (Solarbio, P0100). After centrifugation, the supernatants were boiled for 10 minutes in the SDS-PAGE loading buffer. After being separated by SDS-PAGE at 10%, the proteins were transferred to a PVDF membrane with a thickness of 0.45 μm. The membrane was blocked for two hours at room temperature using 5% skim milk, and then it was incubated overnight at 4 °C with primary antibodies (Proteintech, 22170-1-AP, Zenbio, R381255 and CST, 3558S). After washing the primary antibodies with TBST, the membranes were incubated for 1 hour at room temperature with secondary antibodies (Proteintech, SA00001-2). For viewing, hypersensitive electrochemiluminescence liquid (NCM, P10100) was employed. The bands were analyzed by Image Lab software.

### 2.8 Statistical analysis

All data were expressed as the mean ± SD, and were analyzed by GraphPad Prism. The Shapiro-Wilk test was used to test the normality of the distributions. Unpaired Student-t tests were used for data with normally distributed, while Mann-Whitney U tests were employed for data that depart from normality. Normality distributed data with two variables were evaluated by two-way ANOVA with Sidak’s multiple comparisons. p <0.05 was considered statistically significant.

## 3 Results

### 3.1 Experimental modeling of hypothyroidism in adult mice was established

After six weeks of antithyroid treatment, the hypothyroid mice had a significantly higher serum TSH concentration, while the TT_4_ and TT_3_ concentrations were considerably decreased than the control (Fig. 1A-C). Further, the thyroid gland in hypothyroid mice showed compensatory hypertrophy (Fig. 1D). The body weight of hypothyroid mice was lower than the control as the treatment period was extended (Fig. 1E). In conclusion, we have successfully constructed the hypothyroid model in adult mice by antithyroid treatment.

**Figure 1.**
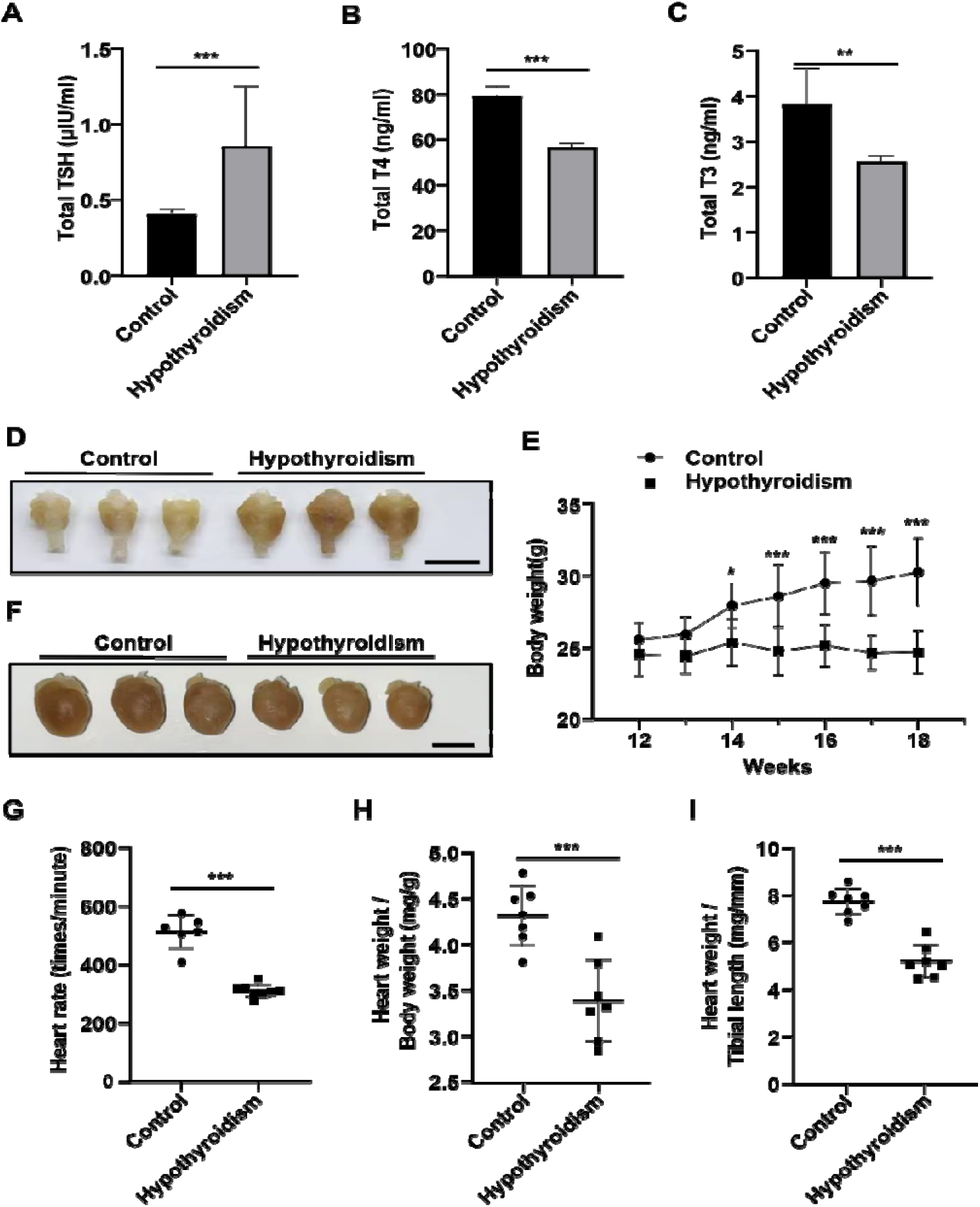
Hypothyroidism induces mouse cardiac atrophy. 12-week-old C57BL/6J mice in the hypothyroid group were provided with drinking water containing 0.1% methimazole and 1% potassium perchlorate for a continuous period of six weeks. **A** Serum levels of TSH were compared to the control (n=7). **B** Serum levels of T4 were compared to the control (n=7). **C** Serum levels of T3 were compared to the control (n=7). **D** Changes in thyroid gland appearance after antithyroid treatment, with the scale bar representing 5 mm. **E** Changes in body weight of mice from week 12 to week 18 (n=7). **F** Changes in heart appearance after antithyroid treatment, with the scale bar representing 5 mm. **G** The heart rate of the mice. **H** Ratio of heart weight to body weight. **I** Ratio of heart weight to tibia length. Data in graphs A and B were analyzed by Mann-Whitney U tests. Data in graphs C and G-I were analyzed by Unpaired Student’s t-tests, with each symbol in graphs G-I representing a mouse. Data in graph E were analyzed using two-way ANOVA with Sidak’s multiple comparisons. (* p<0.05, ** p<0.01, *** p<0.001).

### 3.2 Hypothyroidism induces cardiac atrophy

The heart rate of the hypothyroid mice exhibited a declining tendency (Fig. 1G) when ECG monitoring under anesthesia. This conclusion is in line with the results of a prior study^[11]^. It was observed that the hearts of the hypothyroid mice were smaller than those of the control after the autopsy, (Fig. 1F). Furthermore, the ratios of heart weight to body weight and heart weight to tibial length were both significantly decreased. (Fig. 1H, I).

### 3.3 Quality assessment of the metabolome

LC-MS was used to comprehensively scan 14 individual heart tissue samples from two groups in both POS and NEG ionic modes under ideal conditions. To record signal drift, mixed equal parts of all samples were used as quality control samples (QCs), and inserted at the start, finish, and a few intermediate points during signal data gathering. The QC is clustered in both POS and NEG ionic modes, as seen in Figure 2A, indicating the correction’s good effect. The two groups were more obviously distributed in different regions in the principal component analysis (PCA) score plot. That indicates the metabolite composition structure of the heart in the hypothyroid mice had more obvious differences from the control. The effects of metabolite patterns on the heart after antithyroid drug interventions were examined using the partial least squares discriminant analysis (PLS-DA) and orthogonal projection to latent structure discriminate analysis (OPLS-DA). Figure 2B demonstrates that the PLS-DA model performed better in both ionization modes, and the two groups’ cardiac metabolic profiles exhibit a separation tendency. The results show that the metabolites are remarkably shifted in the mouse heart after antithyroid treatment The difference was further verified by the OPLS-DA, suggesting that the metabolic profile in the heart of the hypothyroid mice was differed from the control. The OPLS-DA R^2^Y and Q^2^ values in each mode were near 1, which indicates the data were without overfitting and were believable (Fig. 2C). The OPLS-DA models’ capacity to accurately discriminate the samples was examined by using 200 permutation tests for cross-validation. It was not overfitted, according to the predictive ability (Q^2^) and goodness-of-fit (R^2^) values (Fig. 2D).

**Figure 2.**
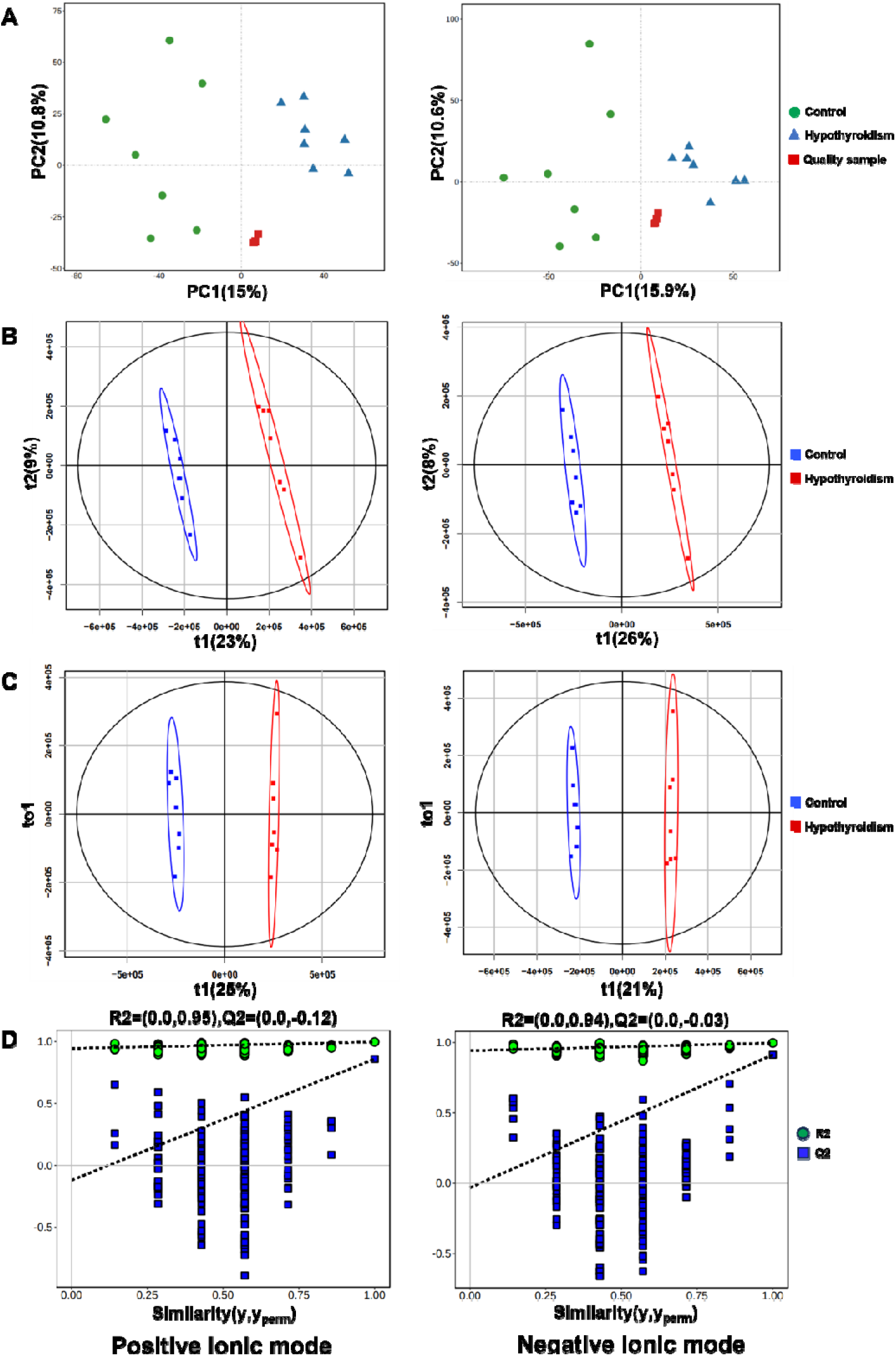
Hypothyroidism affect metabolomics of mice heart. Multivariate statistical analysis of the mouse heart based on LC-MS metabolomics. **A** Principal component analysis (PCA) score plots in positive and negative ion modes. **B** Orthogonal projection to latent structure discriminant analysis (PLS-DA) score plots in positive (R2Y=0.998, Q2=0.907) and negative (R2Y=0.997, Q2=0.92) ion modes. **C** OPLS-DA score plots in positive (R2Y=0.997, Q2=0.913) and negative (R2Y=0.997, Q2=0.86) ion modes. **D** Permutation test for OPLS-DA score plots in positive (R2= (0.0, 0.95), Q2= (0.0, -0.12)) and negative (R2= (0.0,0.94), Q2= (0.0, -0.03)) ion modes. Each symbol represents a sample in graphs A-C.

### 3.4 Metabolic alterations were identified in the hearts of hypothyroid mice compared with controls

Through detection and calculation, a total of 1254 metabolites were identified in the POS ion mode and 896 metabolites in the NEG ion mode. The screening of significant DEMs depended on the two conditions of variable importance for projection (VIP) over 1.0 in the OPLS-DA analysis model, combined with t-test p < 0.05. Based on the above criteria, compared to the control, the heart in hypothyroid mice had 134 metabolites in the POS ion mode, of which 68 were upregulated and 66 were downregulated. In the NEG ion mode, there were 87 metabolites, of which 59 were upregulated and 28 were downregulated (Supplementary table). Volcano plots were used to show differential metabolites between the two groups in both POS and NEG ionic modes. (Fig. 3).

**Figure 3.**
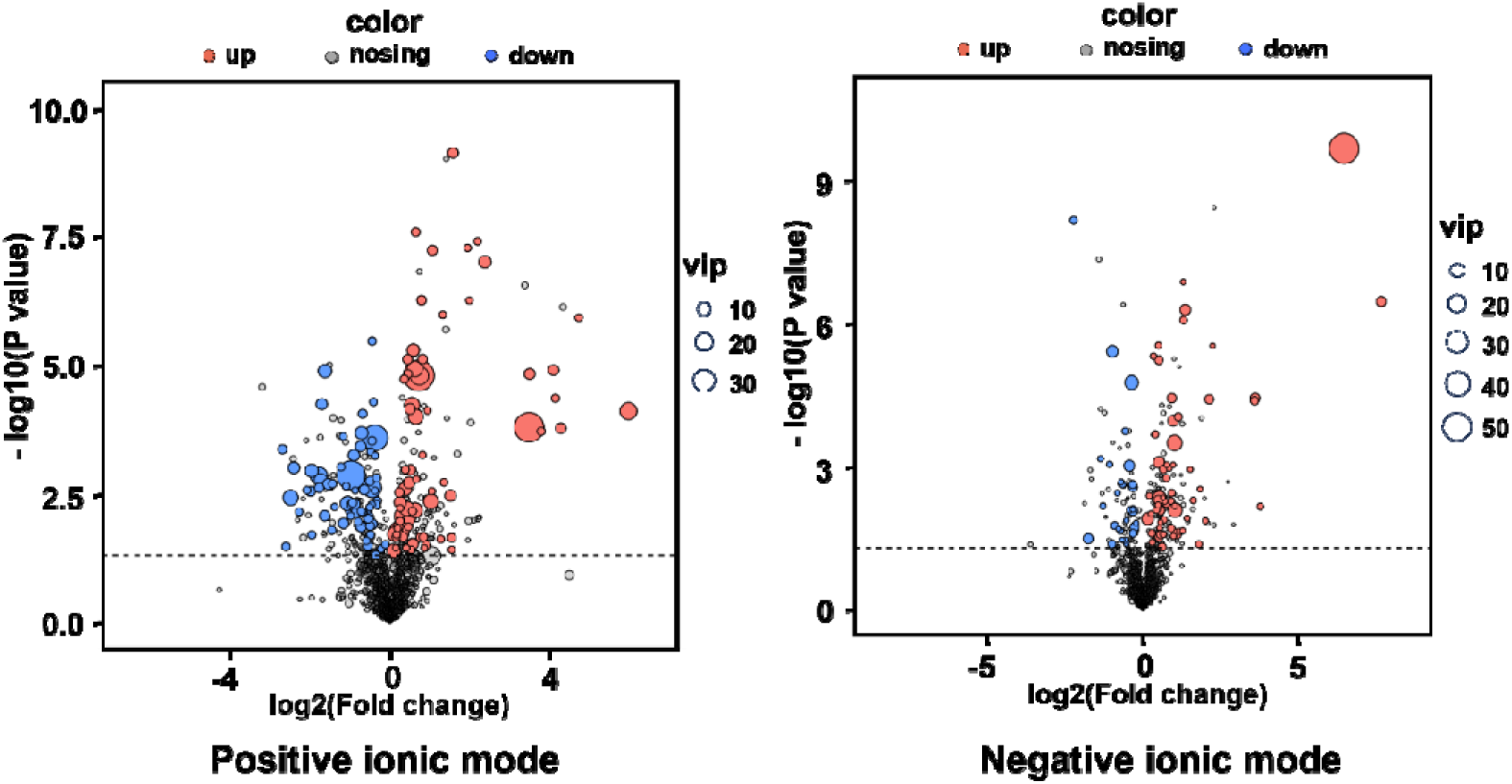
Volcano plots displaying the results of pairwise comparisons in positive and negative ion modes from heart metabolites between hypothyroid mice and control. Each dot represents a metabolite. Red dots indicate metabolites that have been significantly upregulated, blue dots indicate metabolites that have been significantly downregulated, and black dots indicate no significant alteration. The screening criteria are VIP≥1, p<0.05.

In the form of heat maps, an initial illustration of the main categories of DEMs with p <0.05 is provided, such as Phospholipids (Fig. 4), Lipids or Lipid-like, and Carbohydrates and Carbohydrate conjugates (Fig. 5). Notably, the heart in hypothyroid mice had considerably lower levels of seven lipoyl-carnitines, such as Decanoyl-L-carnitine and Lauroyl-L-carnitine and so on, than the control (Fig. 6A). The beta-oxidation of fatty acids provides the mature heart with its primary energy source^[12,13]^. However, the reduction in lipoyl-carnitine is an injury to the heart’s regular energy supply. Additionally, some glycolytic intermediates were found to be elevated, including Fructose-1,6-bisphosphate, D-fructose-1,6-bisphosphate, D-fructose-6-phosphate, and L-(+)-lactic acid (Fig. 6B). Moreover, compounds linked to the PPP, including Ribulose 5-phosphate, D-ribulose-5-phosphate, D-mannose 6-phosphate, 6-phosphogluconic acid, and 6-Phospho-D-gluconate, were also elevated (Fig. 6C). The heart in hypothyroid mice showed markedly enhanced glucose metabolism, as evidenced by the considerable rise in several metabolites from the glycolysis and PPP.

**Figure 4.**
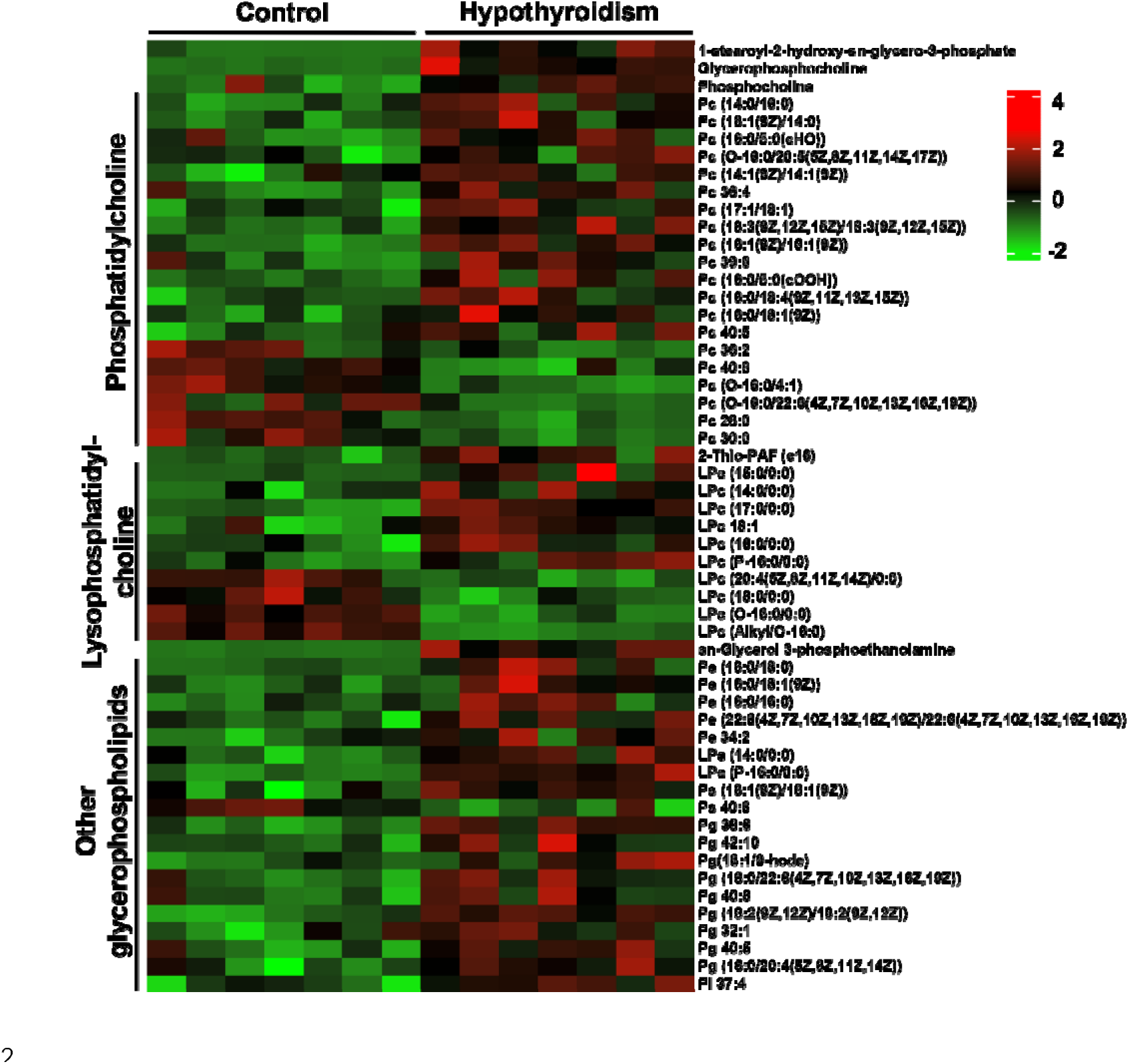
Heat map showing the changes in glycerophospholipid levels of the heart between hypothyroid mice and control. Each metabolite is shown in a row, and each column represents a mouse. Green indicates a relatively low abundance of metabolites, while red indicates a high abundance.

**Figure 5.**
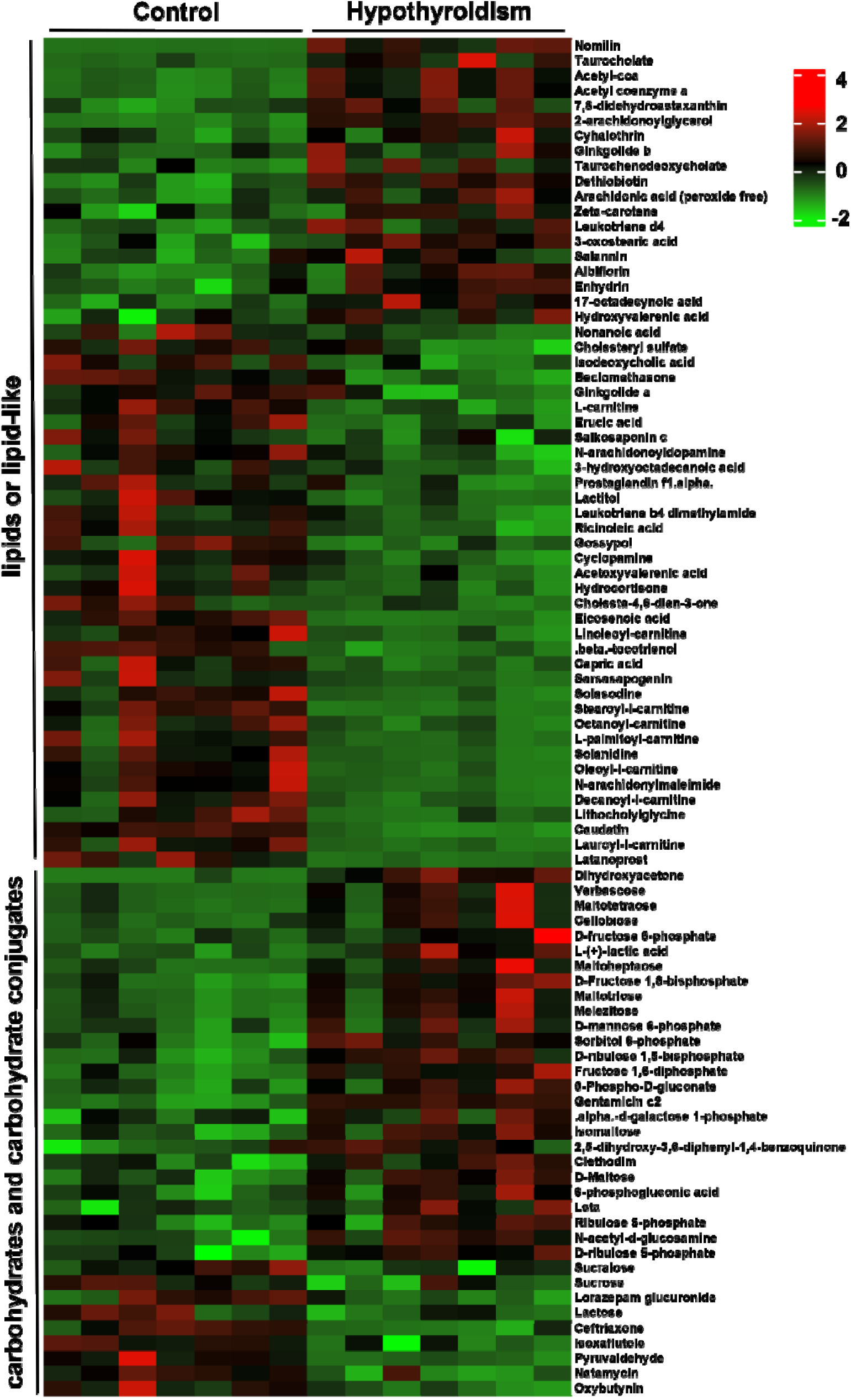
Heat map showing the changes in the lipids or lipid-like molecules, and carbohydrates and carbohydrate conjugates of the heart between hypothyroid mice and control. Each metabolite is shown in a row, and each column represents a mouse. Green indicates a relatively low abundance of metabolites, while red indicates a high abundance.

**Figure 6.**
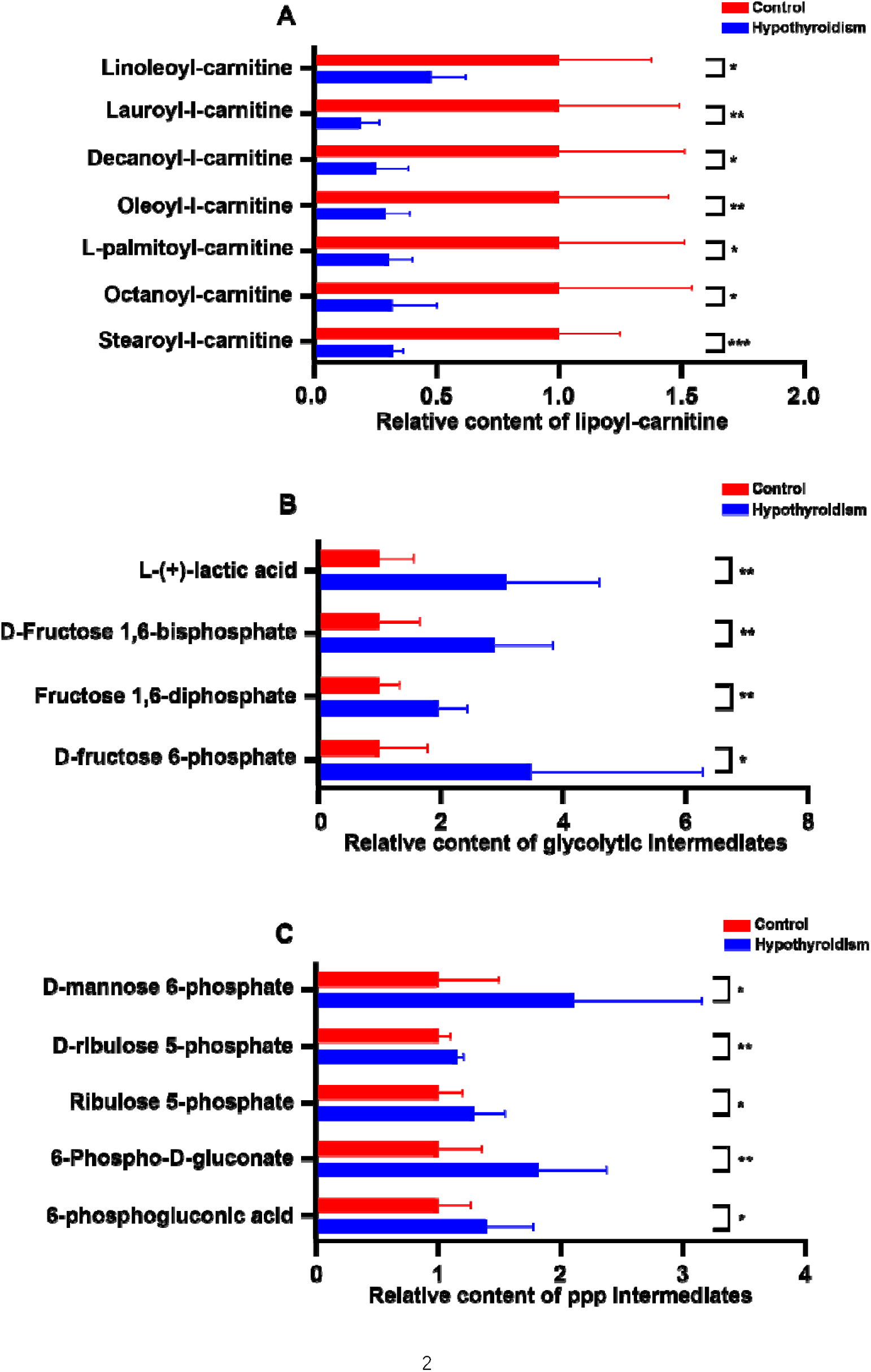
Bar charts representing the significantly altered metabolites in the heart after antithyroid treatment. **A** A dramatic decrease in lipoyl-carnitines occurred in the hearts of hypothyroid mice. **B** Glycolytic intermediates were significantly elevated in the hearts of hypothyroid mice. **C** Intermediates of the pentose phosphate pathway (PPP) were significantly increased in the hearts of hypothyroid mice. (n=7 mice, * p < 0.05, ** p < 0.01, *** p < 0.001).

### 3.5 Hypothyroidism may affect the lipid metabolism pathway in the mouse heart

Referring to the KEGG database, DEMs were mapped into the database for metabolic pathway analysis. Metabolic pathways enriched are shown in Figure 7A, which involve Mitochondrial beta-oxidation of long, medium, and short chain saturated fatty acids, Fatty acid metabolism, Amino sugar metabolism, Citric acid cycle, Glycerolipid metabolism, Phospholipid biosynthesis, and other pathways. To further explore the mechanism of hypothyroidism regulation of metabolic disorders in the heart, the metabolic pathways were analyzed using Metabo Analyst. The results showed that the effect of hypothyroidism on Glycerophospholipid metabolism was more significant (Fig. 7B). These vital metabolic pathways that hypothyroidism may impact are identified, which also offer critical clues for continued investigation of its molecular basis. The distribution and function of DEMs in these pathways can be investigated to better understand how hypothyroidism alters cardiac metabolism and how these alterations impact the structure and function of the heart.

**Figure 7.**
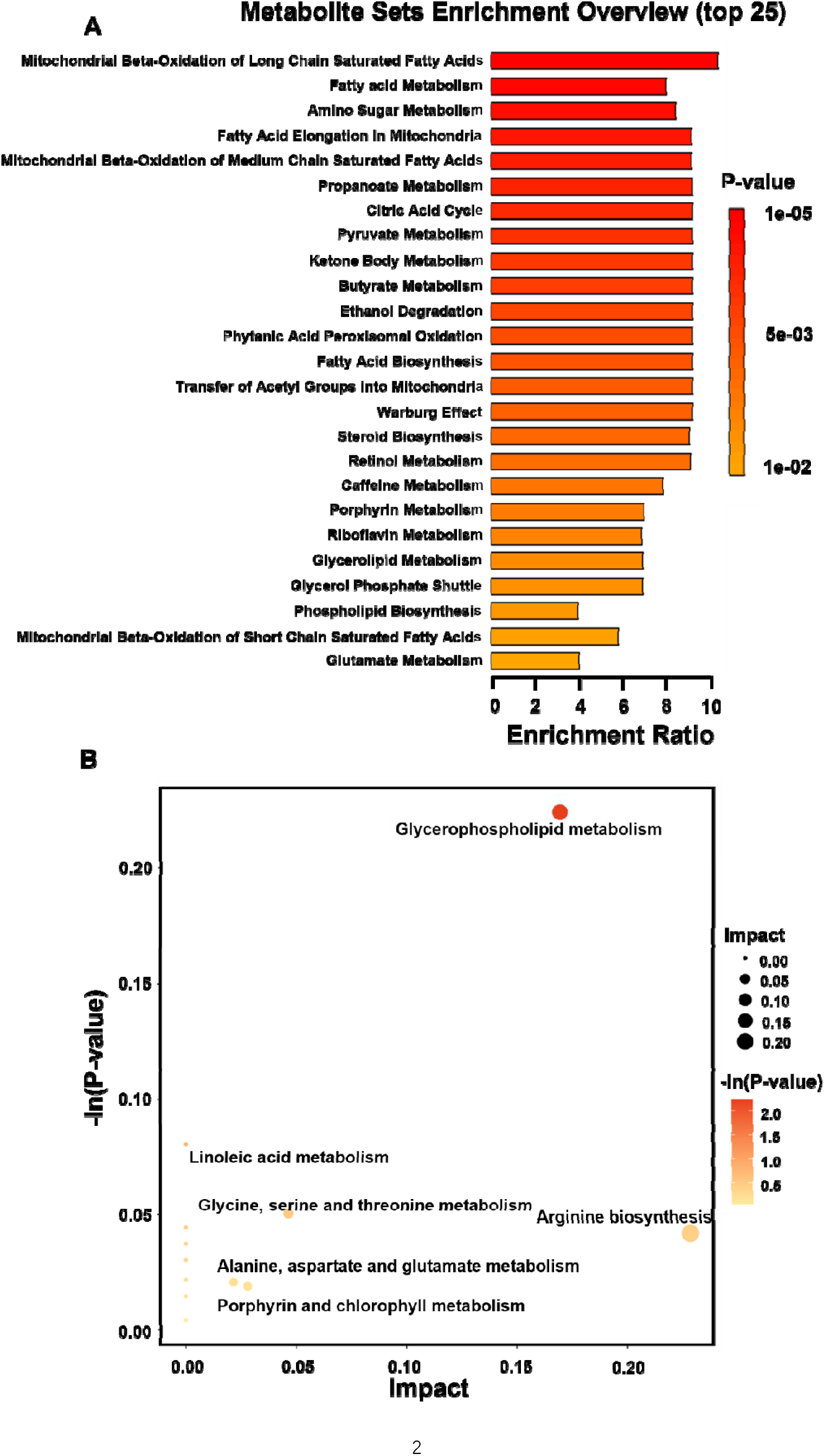
Metabolic pathways involved in hypothyroidism-induced cardiac metabolic disorders. **A** Pathway enrichment analysis of significantly different metabolites based on the KEGG database. **B** Scatterplot displaying the MetaboAnalyst Pathway analysis results. The P-value is represented on a color map in graphs A and B.

### 3.6 Hypothyroidism decrease the protein level of CPT1B and CPT2, and increase the level of LDHA in mouse heart

To further substantiate the reduction in acylcarnitine levels and the increase in glycolytic intermediates. The principal enzymes of fatty acid oxidation, CPT1B and CPT2, exhibited significantly decreased protein expression in the heart of hypothyroid mice compared to the control, as shown in Figures 8A and B. Sametime, we observed a significant elevation in the relative expression of LDHA which important plays a role in glycolysis. (Fig. 8C). These findings may explain the phenomenon that hypothyroid mice exhibited lower levels of lipoyl-carnitine and a threefold accumulation of lactic acid compared to the control.

**Figure 8.**
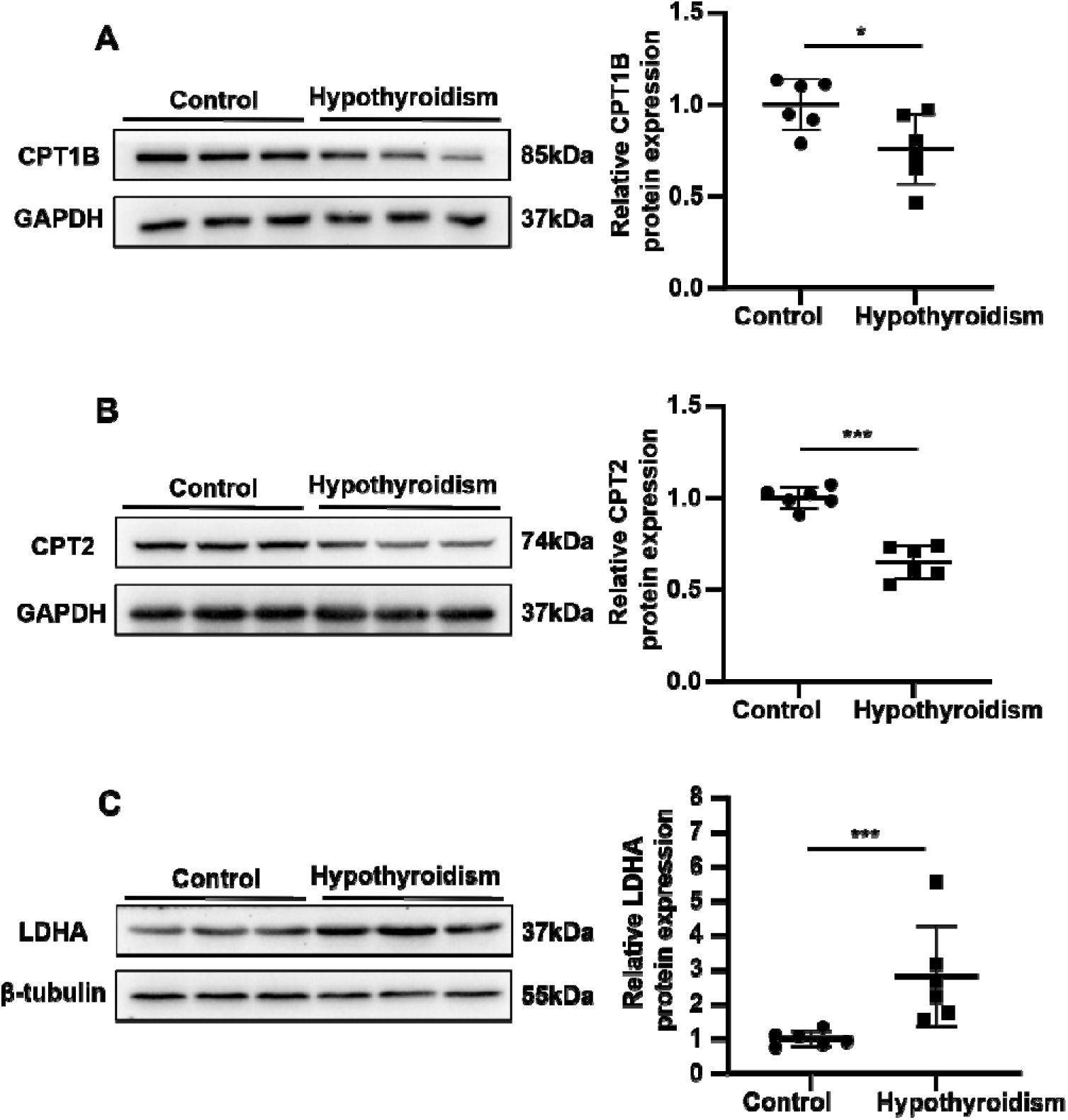
Hypothyroidism decrease the protein level of CPT1B and CPT2, and increase the level of LDHA in mouse heart. 12-week-old C57BL/6J mice in the hypothyroid group were provided with drinking water containing 0.1% methimazole and 1% potassium perchlorate for a continuous period of six weeks. **A** Representative Western blot and semi-quantification statistical data showing the expression of CPT1B protein. **B** Representative Western blot and semi-quantification statistical data showing the expression of CPT2 protein. **C** Representative Western blot and semi-quantification statistical data showing the expression of LDHA protein. Data in graphs A-C were analyzed by Unpaired Student’s t-tests, and each symbol in the right panel represents a mouse. (* p < 0.05, *** p < 0.001).

## 4 Discussion

According to earlier research, hypothyroidism can impact several systems and cause a hypometabolic condition via a wide range of complex mechanisms^[14,15]^. This study established a hypothyroid model in adult mice through antithyroid drug treatment, and observed significant changes in the mice’s hearts, such as reduced heart rate and cardiac atrophy. However, the underlying molecular mechanisms, especially those related to energy metabolism and myocardial function, remain poorly understood. This study pioneers the use of untargeted metabolomics for a comprehensive analysis of cardiac metabolite changes, revealing decreased acylcarnitine levels and increased glycolytic intermediate levels in the hearts of mice with hypothyroidism. These findings provide novel insights into the pathophysiology of the condition. Considering that the heart is one of the most metabolically active organs in the human body^[16]^, a decrease in its metabolic capacity can have a severe impact on cardiac function, as shown by diastolic and systolic dysfunction^[5–7,11]^ as well as a decline in heart rate. We speculate that the above-mentioned symptoms may result from hypothyroidism-induced alterations in cardiac energy metabolism, resulting in the heart with inadequate energy to function properly. Even though the heart can use a variety of energy substrates such as fatty acids, lactate, glucose, and so on to maintain adenosine triphosphate (ATP) production. Furthermore, 40–60% of ATP production is dependent on fatty acid oxidation in healthy cardiomyocytes, with glucose oxidation or glycolysis playing a smaller _role_^[12,13]^.

CPT1B catalyzes the transfer of long-chain fatty acids from acyl-coenzyme A to carnitine, leading to the production of lipoyl-carnitine, the initial step in the flow of long-chain fatty acids into the mitochondria^[17]^. Our findings indicate that hypothyroidism significantly compromises the capacity for fatty acid oxidation in the heart, as evidenced by reduced expression of CPT1B and CPT2 and decreased level of lipoyl-carnitines. In addition, the enrichment of metabolic pathways for the beta-oxidation of long, medium, and short chain saturated fatty acids also provides evidence for this impairment. The availability of fatty acids for mitochondrial beta-oxidation is limited, which is the primary energy source in healthy cardiomyocytes. Furthermore, LDHA was overexpressed and several glycolysis intermediates were markedly elevated in the heart of hypothyroid mice. This suggests that the heart of hypothyroid mice might exhibit an elevated glycolytic capacity compared to the normal. However, the majority of heart diseases are linked to adaptive modifications in energy metabolism, such as a preference for glucose over fatty acids as an energy substrate and a slow decline in mitochondrial oxidative phosphorylation^[12,18]^. These metabolic alterations resemble changes observed in heart failure (HF). Glycolysis is induced as a compensatory response for decreased mitochondrial oxidative phosphorylation and ATP production in HF, but this response is insufficient to restore cardiac function or fully compensate for energy deficits in HF^[19]^. In addition, the heart is burdened by the accumulation of lactate brought on by increased glycolysis^[20]^. Moreover, the PPP, which may play a dual role in regulating redox homeostasis, is activated in HF models, resulting in increased superoxide production, thereby affecting the heart^[21]^. Building on these considerations, cardiac energy metabolism alteration possibly contributes to the development of myocardial injury and cardiac dysfunction in individuals with hypothyroidism.

In the present study, some pathways— such as Mitochondrial beta-oxidation of long, medium, and short chain saturated fatty acids, Fatty acid metabolism, Citric acid cycle, Glycerophospholipid metabolism, Phospholipid biosynthesis— were likely associated with the development of hypothyroidism-induced myocardial atrophy. Among these, one of the most significantly changed pathways was the Glycerophospholipid metabolism pathway. Furthermore, an increased risk of many cardiovascular diseases has been correlated with dysregulated lipid metabolism, specifically with disturbed glycerophospholipid metabolism^[22]^. Numerous physiological processes involve glycerophospholipids, which are abundant phospholipids found in cell membranes. Dyslipidemia and endoplasmic reticulum stress are examples of metabolic disorders that can result from disturbed glycerophospholipid metabolism^[23]^. Glycerophospholipid metabolism was identified as specific to the development of HF by previous untargeted metabolomic analysis of serum from HF patients. They observed that baseline levels of phosphatidylcholine (PC) and lysophosphatidylcholine (LPC) were decreased in patients with HF, resulting in disorders of glycerophospholipid metabolism which contributed to the development of HF^[24]^. According to another study, phosphatidylethanolamine (PE) (12:1e/22:0) and PC (22:4/14:1) may be biomarkers for the onset of HF following myocardial infarction, offering a novel approach to HF prevention^[25]^. Additionally, glycerophospholipid metabolism was found to be the most significantly altered pathway in the plasma metabolomics analysis of various coronary artery disease subgroups in all pairwise comparisons^[26]^.

The study’s findings showed that the hearts of hypothyroid mice had 49 significantly different glycerophospholipids compared to the control (p < 0.05). These included 20 PC, 10 LPC, 5 PE, 2 lysophosphatidylethanolamine (LPE), 2 phosphatidylserine (PS), 9 phosphatidylglycerol (PG), 1 phosphatidylinositol (PI). These imply that hypothyroidism impacts glycerophospholipid metabolism, which in turn may impact the heart and its function. PC is one of the most abundant phospholipids in cardiac cell membranes and is essential for maintaining the integrity and functionality of cell membranes as well as for cell signaling and cytoskeletal stability^[27]^. In vitro and in vivo, LPC may alter the physiology of pericytes, neuronal cells, and the vascular endothelium, indicating that it could be a strong risk factor and is linked to the pathogenesis and prognosis of cardiovascular diseases^[28,29]^. Likewise, PE, PS, PG, and PI are all involved in different biological activities that occur within cells^[29,30]^. In the mouse model, hypothyroidism appears to have caused a disorder in myocardial glycerophospholipid metabolism. However, more thorough research is still needed to determine the precise molecular mechanisms that link this metabolic alteration to hypothyroidism-induced myocardial atrophy.

Thus, the aforementioned findings offer fresh insights into the molecular mechanisms behind hypothyroidism-induced myocardial atrophy, and they could aid in the diagnosis and management of patients with hypothyroid cardiomyopathy. However, these more specific mechanisms need to be verified by additional studies in the future.

## Author Contribution

FL and HG designed and supervised the experiments. HG and WX performed experiments. FL, HG, and SW analyzed the data. The first draft of the manuscript was written by HG and YY. All authors read and approved the final manuscript.

## Acknowledgements

Appreciate the technical support provided by Guangzhou Kidio Biotechnology Co., Ltd.

## Funding

This work was supported by National Natural Science Foundation of China (82160063).

## Declarations

## Conflict of interest

The authors declare that they have no relevant financial or non-financial interests to disclose.

## Ethics approval

All experimental procedures were approved by the Institutional Animal Care and Use Committee of Guilin Medical University.

## Notes

### Competing Interest Statement

The authors have declared no competing interest.

